# Examination of 2D frontal and sagittal markerless motion capture: Implications for 2D and 3D markerless applications

**DOI:** 10.1101/2023.01.17.523930

**Authors:** Logan Wade, Laurie Needham, Murray Evans, Polly McGuigan, Steffi Colyer, Darren Cosker, James Bilzon

## Abstract

This study examined if occluded joint locations from markerless motion capture produced 2D joint angles with reduced accuracy compared to visible joints, and if 2D frontal plane joint angles were usable for practical applications. Fifteen healthy participants performed over-ground walking whilst recorded by fifteen marker-based cameras and two machine vision cameras (frontal and sagittal plane). Repeated measures Bland-Altman analysis illustrated that markerless standard deviation of bias (random differences) for the occluded-side hip and knee joint angles in the sagittal plane were double that of the camera-side (visible) hip and knee. Camera-side sagittal plane knee and hip angles were near or within marker-based error values previously observed. While frontal plane random differences accounted for 35-46% of total range of motion at the hip and knee, systematic and random differences (−4.6-1.6 ± 3.7-4.2°) were actually similar to previously reported marker-based error values. This was not true for the ankle, where random difference (±12°) was still too high for practical applications. Our results add to previous literature, highlighting shortcomings of current pose estimation algorithms and labelled datasets. As such, this paper finishes by reviewing marker-based methods for creating anatomically accurate markerless training data.

## INTRODUCTION

The accuracy and reliability of markerless motion capture for biomechanical applications has been steadily improving over the past two decades Open-source markerless motion capture paired with low-cost cameras, provides researchers, clinicians and coaches with an accessible, cost-effective and transparentmotion capture solution ^4-9^. However, determining the accuracy of markerless systems is difficult due to scarcity and complexity of gold standard methods such as biplanar radiographic imaging ^10,11^ and videoradiography ^12,13^. Instead, comparisons are often performed against marker-based systems, although, inherent marker-based errors such as soft tissue artefact^13,14^, marker placement unreliability ^15,16^ and altered participant movement patterns ^17^ will obscure the markerless results.

2D markerless motion capture from a single camera view could facilitate analysis of clinical and sporting gait parameters, without the need for specialist personnel to process the data ^5,18^. Furthermore, future pose estimation algorithms could be employed to extract additional key variables from large clinical video databases that have been acquired over many years ^5,18^. Previous work has examined sagittal plane kinematics during walking and underwater running ^4,6^, identifying joint angles that are in line with those obtained using manually labelled methods. However, only angles of joints nearest to the camera were examined, despite pose estimation algorithms also providing occluded joint centre locations on the far side of the body ^19^. Current open access pose estimation training datasets have labelled both occluded and visible key points (joint centre locations) ^20,21^, which is useful for entertainment applications, ensuring that detected body parts do not simply disappear when occluded by other limbs. However, from a biomechanical point of view, manually labelling occluded joints requires the labeller to estimate where the joint centre is located, likely resulting in an inevitable reduction in accuracy and reliability. While occluded joints may have reduced accuracy compared to visible joints, the difference could potentially be small enough to have a minimal effect for some applications, especially in only partially occluded joints (e.g. ankle and knee during gait). However, to date, there has been no comparison of the accuracy and reliability of joint angles obtained from occluded and visible joints, when using 2D open-source pose estimation algorithms.

In the frontal plane, joint occlusion is minimal, however limited movement requires high precision and low random differences to detect small but meaningful changes. Both Shin, et al. ^18^ and Azhand, et al. ^7^ have examined walking gait in the frontal plane, with Azhand, et al. ^7^ notably collecting their data using a handheld smartphone camera. However, both studies only measured temporospatial measures and did not examine joint angles. As such, currently there has been no evaluation of frontal plane joint angles using 2D markerless methods, leading to uncertainty of usability for clinical and sporting applications.

This paper aims to explore two interconnected questions by comparing open-source single camera 2D pose estimation (OpenPose) with marker-based motion capture. Firstly, are occluded joint angles and joint centre locations identified with reduced accuracy compared to visible joints? Secondly, are markerless frontal plane joint angles sufficiently accurate for practical applications? It was hypothesised that occluded joint angles in the sagittal view would be too variable for biomechanical applications. It was also hypothesised that markerless motion capture will likely be too variable to detect meaningful changes in frontal plane joint angles.

In this study, fifteen healthy participants performed over-ground walking at their preferred speed, while being recorded by fifteen marker-based cameras and two machine vision cameras (frontal and sagittal plane). The OpenPose ^19^ pose estimation algorithm was used to identify joint centre locations in each machine vision camera view. 3D marker-based motion capture data were reprojected into each 2D machine vision camera plane. Comparison between methods were performed using repeated measures Bland-Altman analysis ^22^, which ensured that random differences were not underestimated. Systematic difference (bias) and random difference (standard deviation of bias) of the joint angles and joint centre locations in the frontal and sagittal plane were compared between 2D markerless motion capture and reprojected 3D marker-based motion capture. To clearly describe differences in the sagittal plane, joint angles and locations in this plane will henceforth be referred to as camera-side (right side joints) and occluded-side (left side joints).

## Results

### Sagittal Plane Joint Centre Location

On average, systematic differences for the camera-side and occluded-side joint centre locations of the ankle and knee were similar (8.8-12.0 pixels), while occluded-side hip and shoulder joint centre locations (11.6-15.2 pixels) were worse than their camera-side counterparts (8.5-8.6 pixels, Figure 1A). For both sides of the body, systematic and random differences in the ankle and knee had substantial spikes during push-off and swing phase (Figure 1C and 1D), while the hip and shoulder joint centre locations were relatively constant over the stride (Figure 1A, 1E and 1F). Systematic and random difference of the metatarsophalangeal (MTP) joint for both the camera (20 ± 9 pixels) and occluded-side (23 ± 10 pixels) were worse than all other joints (Figure 1A), which was produced primarily during the push-off and swing phase of walking (Figure 1B). Sagittal plane, repeated measures Bland-Altman results for joint centre locations can be found in Table 1 of Supplementary File 1.

**Table 1:** Repeated measures Bland-Altman analysis of the 2D markerless ankle, knee and hip joint angles in the sagittal and frontal plane, relative to reprojected marker-based motion capture. Camera-side represents the right side of the body, which was closest to the camera, while the occluded-side represents the left side of the body, which was furthest from the camera in the sagittal view. Left and right sides of the body were combined in the frontal plane.

| Joint Angles |  | Bias (°) | Standard Deviation of Bias (°) | LOA (°) |
| --- | --- | --- | --- | --- |
| Sagittal Plane |  |  |  |  |
| Ankle | Camera-side | -8.4 | 5.2 | -18.6 – 1.9 |
|  | Occluded-side | -9.3 | 7.3 | -23.6 – 5.1 |
| Knee | Camera-side | 1.5 | 4.1 | -6.5 – 9.6 |
|  | Occluded-side | 1.6 | 6.9 | -12.0 – 15.2 |
| Hip | Camera-side | -3.6 | 4.6 | -12.6 – 5.3 |
|  | Occluded-side | -4.6 | 9.5 | -23.2 – 14.0 |
| Frontal Plane |  |  |  |  |
| Ankle |  | 0.2 | 12.0 | -23.4 – 23.8 |
| Knee |  | 1.6 | 4.2 | -6.7 – 9.9 |
| Hip |  | -4.6 | 3.7 | -11.5 – 3.0 |

**Figure 1:**
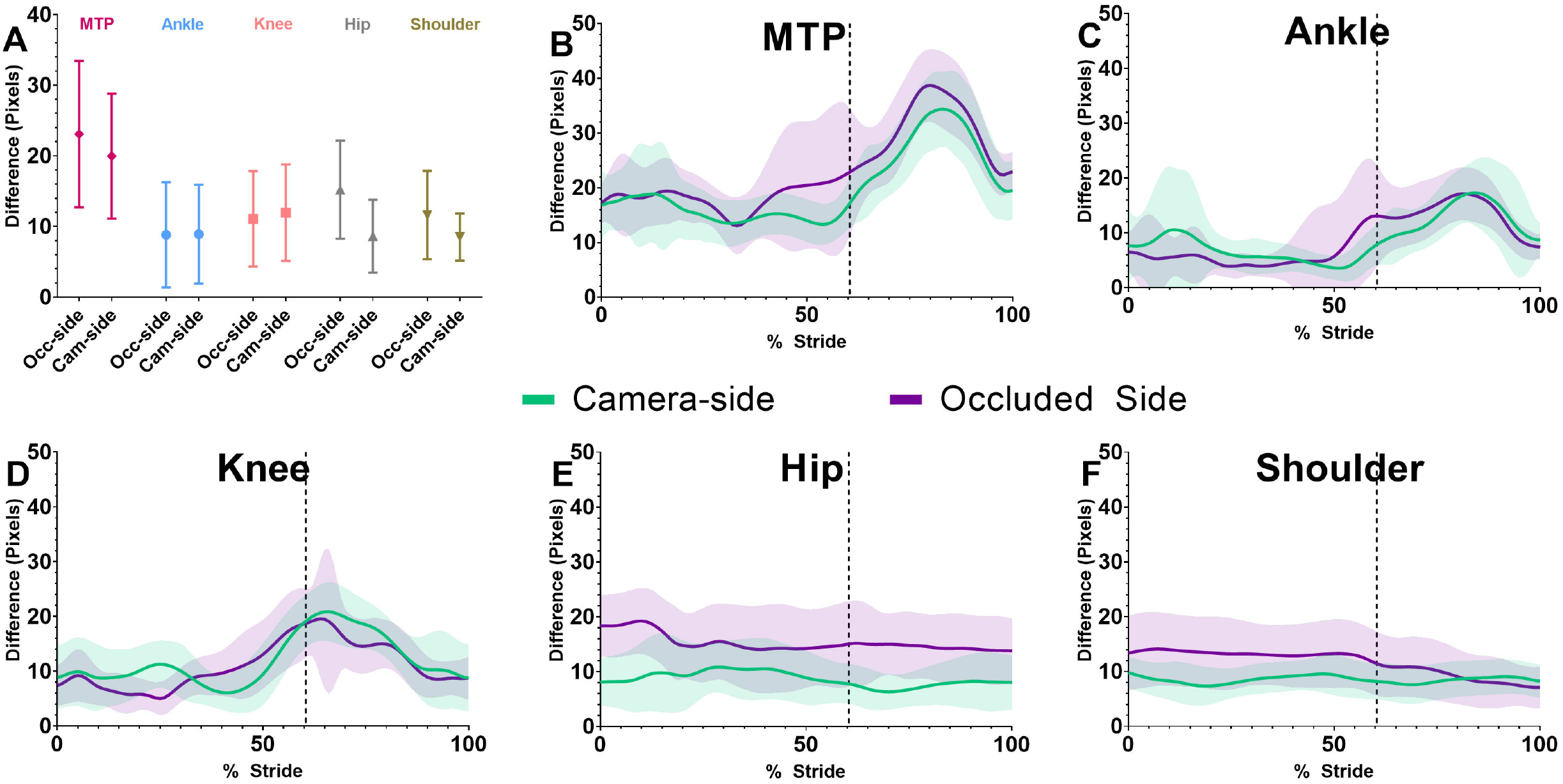
Sagittal plane joint centre location differences over the whole stride (A) for the camera-side (Cam-side) and occluded-side (Occ-side), created using repeated measures Bland-Altman bias (systematic difference) and standard deviation of bias (random difference). In Figure 1B-F, frontal plane joint centre locations were time normalised to one stride and then, within stride mean and standard deviation differences were calculated between markerless and marker-based motion capture for each joint, averaged over all trials. Vertical lines in Figures 1B-F represents the end of the stance phase (60.4% of stride).

### Sagittal Plane Joint Angle

Sagittal plane systematic differences (bias) across the ankle, knee and hip were relatively consistent between camera and occluded sides (slightly higher for the occluded side). However, absolute random differences (standard deviation of bias) were 40%, 68% and 107% greater in the occluded ankle, knee and hip respectively, compared to their camera-side counterparts (Table 1, Figure 2A-C). Average joint angle range of motion (RoM) obtained from the marker-based method was 37 ± 6 °, 45 ± 6° and 67 ± 4° for the ankle, knee and hip respectively. As such, markerless random difference accounted for 14% and 20% of total ankle joint RoM on the camera-side and occluded-side joint angles respectively. Alternatively, in the knee and hip, markerless random difference accounted for 7-9% RoM in camera-side joints and 14-15% RoM in occluded-side joints. Timeseries data of sagittal plane joint angles and differences are presented in Supplementary File 1.

**Figure 2:**
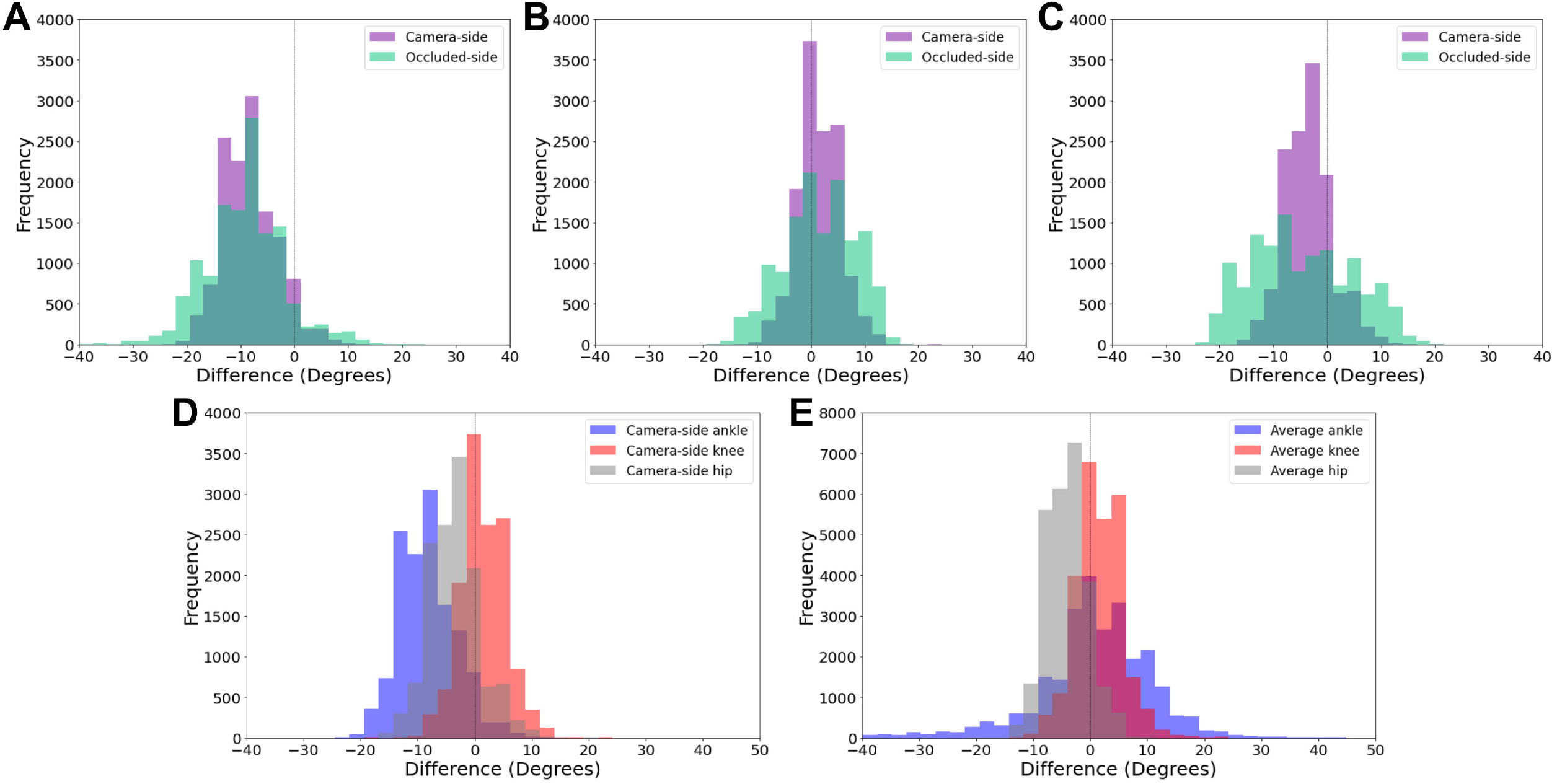
Distribution of joint angle differences between markerless and marker-based methods, for all individual data points (101 timepoints x successful trials x participants). Figures 2A-C illustrate the differences between camera-side and occluded side joints of the ankle (A), knee (B) and hip (C). Figure 2D illustrates the camera-side only differences for the ankle, knee and hip. Figure 2E illustrates the differences for the combined left and right ankle, knee and hip in the frontal plane.

### Frontal Plane Joint Centre Location

Systematic differences of 2D joint centre locations were relatively consistent across all joints (Figure 3A), lowest at the ankle (7.6 pixels) and highest at the shoulder (11.9 pixels). However, random difference (standard deviation of bias) of joint centre locations (Figure 3A) was greatest in the MTP (9.6 pixels) and decreased as joint level increased, smallest at the shoulder (4.8 pixels). Examination of systematic and random differences relative to marker-based motion capture (Figure 3B) indicated that there was a large increase after push-off in the MTP, ankle and knee joint centre locations (Figure 3B), while the hip and shoulder were relatively similar throughout the stride. Frontal plane repeated measures Bland-Altman table for joint centre locations can be found in Table 1 of Supplementary File 1.

**Figure 3:**
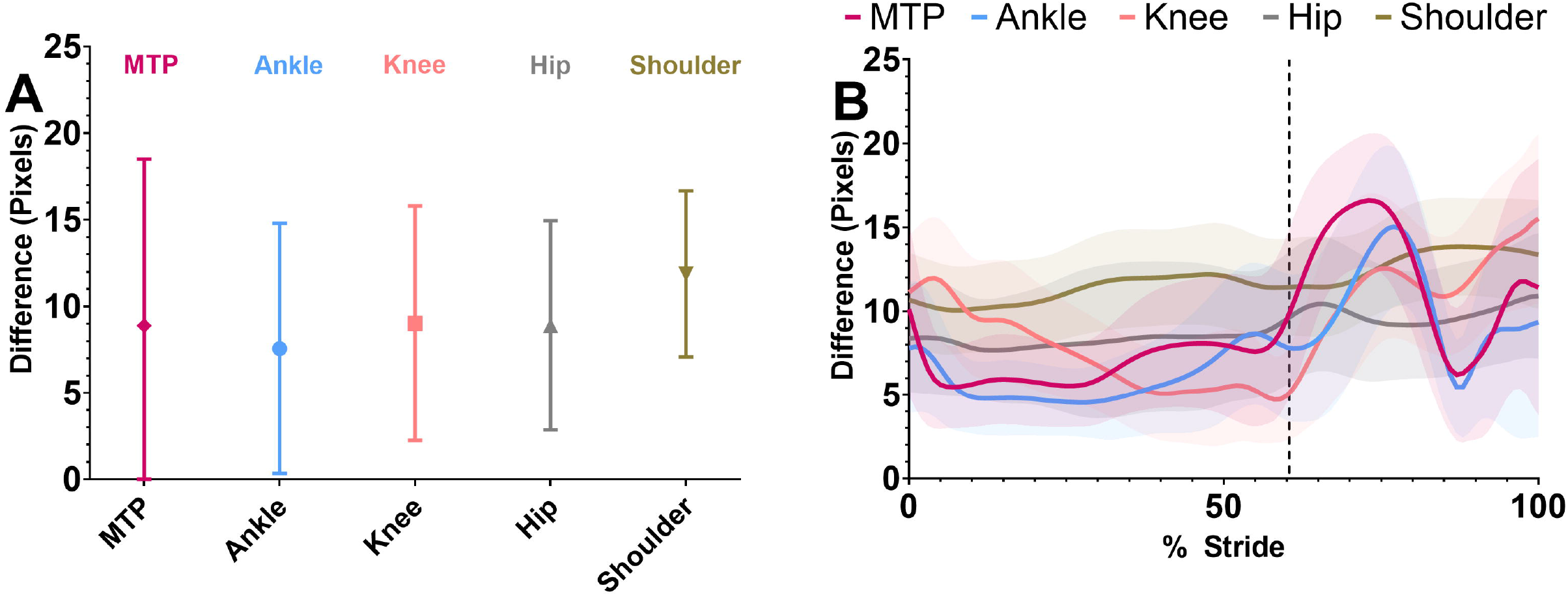
Frontal plane joint centre locations over the whole stride (A), calculated using repeated measures Bland-Altman bias (systematic difference) and standard deviation of bias (random difference). Frontal plane joint centre locations were time normalised and within stride mean and standard deviation differences were calculated (B) between markerless and marker-based motion capture for each joint averaged over all trials. Vertical lines in Figures 3B represent end of the stance phase (toe-off).

### Frontal Plane Joint Angle

Systematic and random differences were relatively similar in the frontal plane compared with sagittal plane about the knee and hip (Table 1). However, while systematic differences were improved about the ankle angle relative to the sagittal plane, random differences were substantially increased (Table 1, Figure 2E). Mean total joint RoM from marker-based data was 31 ± 15° for the ankle, 12 ± 5° for the knee and 8 ± 3 ° for the hip. Due to the small RoM at the hip and the knee, and the large random difference at the ankle, frontal plane markerless random difference accounted for 39%, 35% and 46% of the total RoM about the ankle, knee and hip respectively. Timeseries data of frontal plane joint angles and differences are presented in Supplementary File 1.

## Discussion

Sagittal plane random differences of occluded joint angles were higher in more proximal joints, with the hip being highest (Table 1, Figure 2). Because occluded-side hip and shoulder joint centre location were obstructed for the entire stride, they were more affected than the MTP, ankle and knee (Figure 1). Therefore, the occluded-side knee angle had lower random difference as it was calculated using vector angles between the ankle, knee and hip joint centre locations, compared to the hip angle, which used the knee, hip and shoulder joint centre locations. The knee and hip ratio of random difference to total joint RoM was relatively small in the camera-side joints (7-9%), while the occluded-side knee and hip joints were almost double (15%). Alternatively at the ankle, the random difference was only slightly higher in the occluded-side compared to the camera side (Table 1). Overall, the ankle ratio of random difference to total joint RoM was higher than the knee and hip, which was likely due to two causes. Firstly, the RoM about the ankle is much lower than the knee and hip, and thus the relative proportion of error to signal is lower. Secondly, the markerless system identifies the end of the foot by labelling the hallux (first digit) and fifth digit of the foot, while marker-based motion capture uses the first and fifth MTP joint. Therefore, markerless ankle joint angle will be influenced by toe extension during push-off ^23^. These results indicate that occluded hip and knee joints have roughly double the variability of the camera-side joints, and overall the ankle angle has considerable inaccuracies due to biomechanically limited labelling of the foot in the COCO open access training dataset ^21^ used by the OpenPose algorithm ^19^ we employed.

Frontal plane joint angle systematic differences were relatively small for the ankle and knee, while the hip exhibited a larger offset (Figure 2), although this is most likely due to the systematic differences previously identified in this markerless training dataset ^24^. The ankle angle had considerably worse random differences than the hip and the knee (Figure 2E and Table 1), which was due to the MTP, ankle and knee joint centre locations having increased differences during push-off, swing phase and heel-strike (Figure 3), when knee flexion, dorsiflexion and toe extension may produce occlusion of the MTP and ankle joint.

These results highlight several implications of open-source 2D markerless motion capture (OpenPose) for clinical and sporting applications. Firstly, random differences were almost double in occluded-joint angles compared to visible camera-side joints, and therefore, occluded joints should be discounted when practitioners and coaches are assessing results. Secondly, both occluded and camera-side MTP, ankle and knee joint centres had increased systematic differences during the swing phase, thus, these periods may currently need to be omitted to facilitate examination of joint angles. Thirdly, the MTP joint definition from OpenPose does not account for toe flexion during push-off, which further reduces the accuracy of the ankle joint angle ^23^. Finally, in 2006 Benoit, et al. ^25^ used bone-pin markers to examine the knee joint angle error rate of skin markers during walking, finding marker-based knee flexion/extension errors of 2.8 ± 2.6° and knee abduction/adduction errors of 4.4 ± 3.2°. In comparison, our markerless knee angle differences in the sagittal plane was 1.5 ± 4.1° (flexion/extension), and in the frontal plane was 1.6 ± 4.2° (adduction/abduction), thus markerless differences compared to marker-based motion capture are actually near or within marker-based error rates for the knee in the sagittal (camera-side only) and frontal plane. Benoit, et al. ^25^ obtained absolute errors values at foot-strike, mid-stance and toe-off, while our study examined the entire stride, finding higher variation of joint centre locations during the swing phase (Figure 1). Therefore, these differences would likely reduce further if we only examined the stance phase as Benoit, et al. ^25^ did. Due to similar RoM and differences for the knee and hip angle in our study, these justifications likely extend to the hip as well. Unfortunately, ankle differences were still too high when compared to a study that examined ankle marker-based errors against biplanar fluoroscopy^13^. The authors observed ankle plantarflexion errors ranging maximally up to 4.2° and frontal plane maximal ranging maximally up to 6.3°, while our study observed 95% LOA ranging up to 18.6° in the sagittal plane and 23.8° in the frontal plane. The use of marker-based methods is widespread and often accepted as a gold standard, despite these known errors. These results reinforce the need for comparisons of markerless methods against gold standard measures (e.g. bone pins and biplanar fluoroscopy) to truly determine the accuracy and reliability of markerless systems. However, our findings suggest that during walking, 2D sagittal and frontal plane kinematics identified by the OpenPose algorithm could be obtained for the visible knee and hip joint angles during stance, while the sagittal ankle angle is likely only usable during early and mid-stance and frontal ankle angle is too variable.

The results from this study also have implications for 3D motion capture, due to observed difference between occluded and visible (camera-side) joint centre locations. Manual labelling within pose estimation training datasets requires human labellers to estimate the location of occluded joints, thus it is not surprising that pose estimation algorithms will also have increased uncertainty for occluded joints. Occluded joints will therefore increase the random error and variability of the 3D fusion solution from multiple 2D camera views, especially compared to a system that only performs fusion using joint centre locations of visible joints. Unfortunately, open-source pose estimation algorithms such as OpenPose ^19^ generally do not appear to differentiate between visible and occluded joints, despite some datasets providing this information^21,26^. Additionally, Cronin ^27^ suggested that the inclusion of indiscriminate occluded joints when training the pose estimation algorithm might also reduce the accuracy of detecting all joints, due to the pose estimation algorithm learning that a joint centre location may appear on a point of the body that is not the actual joint. Assessing this theory requires comparison of two identical pose estimation algorithms that differ only in their training dataset (i.e., trained with and without occluded joints), which was beyond the scope of this paper. As such, creation of anatomically accurate markerless datasets should seek to label and classify both occluded and visible keypoints ^21,26^, which may facilitate creation of highly accurate training datasets, while also making these datasets functional for broader applications (e.g. virtual reality).

A potential alternative to 2D single camera markerless systems are minimal 3D systems, which could provide an open-source motion capture system that is cheap, fast, has low space requirements, and minimises parallax and perspective errors. Such systems could perform optimisation of joint centre locations from two or three cameras, which could help to reduce variability. One such open-source system that has recently been released is OpenCap^8^, which combines two or more IOS devices with cloud-computing and OpenSim^28^. Additionally, OpenCap have taken the 20 keypoint outputs (joint centre locations) from OpenPose^19^, and used them as inputs for a second neural network which then infers the location of 43 virtual markers (three virtual markers per segment) and also adds temporal consistency. In their validation paper, OpenCap demonstrated that their double neural network could perform pose estimation and biomechanical modelling with improved kinematic accuracy compared to if pose estimation was used alone^8^. However, improvements were primarily produced in the pelvis and lumbar spine, with minimal or no improvement at the hip, knee and ankle. Additionally, the mean absolute error and root mean squared error for their joint angles appear to have been obtained by performing multiple averaging of their results (averaged across trials and then averaged across participants), which means the root mean square error they presented will likely be underestimated (Supplementary File 2). As such, the root means square error of their ankle, knee and hip angle would likely be closer to the random differences observed by our 2D single camera method, as both systems employed the OpenPose pose estimation algorithm ^19^. While, current minimal 3D camera systems may not produce any improved accuracy over a single 2D camera system for visible joints, system setup is less restricted as it can obtain both frontal and sagittal plane angles in a single trial with minimal occlusion.

Our research supports previous papers ^3,23,24,27,29^ highlighting the need of anatomically accurate training datasets that are fully utilised to enable widespread biomechanical applications. Pose estimation algorithms trained on manually labelled datasets have the potential to outperform marker-based methods, through elimination of soft tissue artefact and inter-session variability ^16^. However, to achieve this, manually labelled image datasets need to have additional key points (3 points on each segment) ^24^, employ labellers with anatomical knowledge ^3,30,31^, and include additional detail (e.g. classifying points as occluded or visible). Unfortunately, the cost and time required to manually label datasets is highly prohibitive, as such, alternative methods for creating datasets are being explored. A potential area of research employs marker-based motion capture to obtain anatomically accurate joint and keypoint locations of interest. To date, there has been no critical review of methods to create labelled datasets using marker-based data, which is key to informing best practices moving forwards. Before we explore this area in more detail, it should be noted that any pose estimation method trained on marker-based data will have known marker-based errors ^13,14,25^ added to the markerless solution, such as marker placement error ^15^ and soft-tissue artefact ^15,17^. Thus, while the creation of datasets using marker-based motion capture could be cheaper, faster and more flexible to advances in machine learning such as volumetric modelling ^32,33^, we need to carefully consider how close markerless systems should match marker-based results.

One method to create labelled 2D images is by recording marker-based and regular video data simultaneously, and then reprojecting marker-based keypoints into the video camera views. However, there are two major drawbacks of this method. Firstly, the pose estimation algorithm will be trained on images of markered participants, and therefore, the algorithm may generalise poorly to new video images that do not include markers. Secondly, marker-based data is generally collected in highly constrained laboratories and therefore the environments and movements within the new training dataset may be limited. Vafadar, et al. ^11^ employed a pose estimation algorithm ^34^ that was first trained on the marker-based Human 3.6M open access dataset ^20^, which was then refined using transfer learning on the ENSAM clinical dataset ^11^. The ENSAM dataset is of particular interest as it combines 3D marker-based motion capture, markerless video and biplanar x-ray (EOS), to ensuring accurate identification of joint centre locations ^11,35^. Vafadar, et al. ^35^ observed significantly improved results after refinement, with considerably decreased systematic differences. However, Human 3.6M ^20^, ENSAM ^11^ and the evaluation performed by Vafadar, et al. ^35^ all employed marker-based motion capture in highly controlled laboratory environments. Thus, their paper likely had a bias of markerless joint angles towards marker-based results, with additional uncertainty surrounding generalisability to video images without markers. Furthermore, they performed a standard Bland-Altman analysis and did not describe how they adjusted their data to deal with repeated measures (Supplementary File 2).

A potentially more promising solution is to create realistic synthetic images ^33,36,37^ using virtual environments and body models from which 2D labelled images can be generated ^33,38^. As a case study, Mundt, et al. ^36^ used marker-based data to generate synthetic images of virtual human models performing a sidestepping task in three participants. While no formal analysis was performed, they did demonstrate similar markerless results between real and synthetic images, although this was only performed for the lower limbs with a blank grey background. Creation of synthetic images using marker-based data must contend with previously described marker-based and markerless errors, in addition domain gap errors ^33^, which are caused by generated images not looking completely real ^39^ or not accounting for the full range of human movement (e.g. hinge joint at the knee in the model). The AGORA training and evaluation dataset has been developed specifically for this purpose, enabling generation of images with diverse movements, environments and cameras angles ^33^, although it does not include biomechanically driven human body models. Future work is needed to determine the biomechanical accuracy of these synthetic images, especially in the clinical and sporting field where movements are so diverse. However, the field of synthetic images is still very novel and therefore, it will likely play a significant role in the development of future markerless motion capture training datasets.

## Conclusion

Our results add to the growing body of literature highlighting potential errors associated with open-source pose estimation datasets and algorithms such as OpenPose, demonstrating a need to discriminate between occluded and un-occluded joints. Sagittal (camera-side) and frontal plane joint angles of the knee and hip produced results that are near or within previously described 3D marker-based motion capture error ranges, although ideally measures should only be taken during stance. The sagittal plane ankle angle may be useable during stance of walking, however the frontal plane ankle angle is not. Minimal 3D markerless motion capture systems may help to reduce occluded joints and loosen data setup restrictions common to 2D motion capture. Given the known problems with marker-based motion capture, care must be taken before employing them for both training and evaluating markerless motion capture systems.

## Methods

Fifteen healthy participants (7 males and 8 females, 26 ± 5 years old, 173 ± 11 cm, 73 ± 14 kg) gave written informed consent to perform overground constant speed walking, while markerless and marker-based motion capture data were recorded. Ethics were approved by the Research Ethics Approval Committee for Health at the University of Bath (EP1819052). Data collection was performed as part of a larger dataset that recorded synchronised 3D marker-based motion capture and 3D markerless motion capture, described in depth by Needham, et al. ^24^. In this study, motion capture data (200 Hz) from fifteen Qualisys cameras (Oqus, Qualysis, Gothenburg, Sweden) and two machine vision cameras (JAI sp5000c, JAI ltd, Denmark) were analysed. One machine vision camera was set up in the sagittal plane (right hand side of the body during walking) and one was set up in the frontal plane (directly in front of the participant). Video data from both systems were time synchronised using TTL pulses sent out from the markerless system. Euclidean spaces of both systems were aligned by moving a single marker through both camera spaces to match corresponding marker locations ^24^. 3D marker-based motion capture data was collected using a full-body marker-set of retroreflective markers attached to the participants upper and lower limbs, pelvis and torso, described in detail by Needham, et al. ^24^.

2D markerless motion capture video data were processed using the open-source pose estimation algorithm OpenPose^19^, which identified 25 keypoints on the participant. In depth implementation of OpenPose on this dataset has been described previously ^24^, however the tracking method used to identify the participant within and across multiple images has not previously been described. While OpenPose is capable of multi-person detection in a single image, it does not associate or ‘track’ people between frames, therefore the person ID given to each individual within an image may change frame to frame. To track only our participant and ignore additional people or false detections, we make use of a classic and highly efficient Kalman filter and the Hungarian algorithm approach ^40,41^. Each detected person’s keypoints are reduced to a single median point and represented by a linear, constant-velocity motion model and confidence value. The confidence threshold was set to 60%, below this value often represented false positive detections, caused by tripods in the background. The motion model, based on a Kalman filter, is then used to propagate a person’s identity in the next frame. The state of the model can be represented as

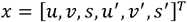

Where u and v represent 2D pixel coordinate locations on the image plane, s is the scale of the detected persons and u’, v’ and s’ represent the rate of change of these respective values between frames. When the detection is associated to a currently tracked person, points u and v are used to update the target state where the velocity components are solved optimally within the Kalman filter. To assign detections to existing persons, each person’s median point is estimated by predicting the new location in the current frame. The assignment cost matrix is then computed as the Euclidean distance between each detection and all the predicted locations of the existing persons. The assignments are then optimally solved using the Hungarian algorithm ^42^. Once all frames had been processed, the participant’s tracked data were collated within a single ID that could be identified from the list of tracked persons, based on the tracked ID with the greatest cumulative distance in pixels, due to the participant being the only person moving substantially during a trial.

Markerless 2D sagittal and frontal plane joint centre locations over the entire trial were smoothed using a bi-directional Kalman filter, which accounts for previous and future locations of each marker. This filtering method has previously been shown to be more effective on markerless data than traditional biomechanical low pass filtering ^24^. Markerless shoulder, hip, knee and ankle joint centre locations were obtained directly from the pose estimation algorithm output, while the MTP joint was defined as the midpoint between the first and fifth toe keypoints. Marker-based data were collected and processed in Qualisys Track Manager (Qualysis, Gothenburg, Sweden) before being exported to Visual3D (C-Motion Inc, Germantown, USA). Hip joint centres were computed via regression from the anterior and posterior superior iliac spine markers as per Visual3D guidelines, knee and ankle joint centres were computed as the midpoint between the medial and lateral markers, shoulder joint centres were offset from the location of shoulder markers using Visual3D guidelines, and the MTP joint centres were computed as the midpoint between the first and fifth MTP joint markers.

Because markerless and marker-based cameras generally cannot occupy the same physical space, previous research has compared results between systems using two separate cameras placed next to each other, resulting in only an approximate alignment between the planes of each system’s Euclidean spaces ^43,44^. This will likely result in some parallax error between cameras that may manifest as systematic differences between the two systems. To mitigate this issue and provide a high-quality comparison between markerless and marker-based results, 3D marker-based joint centre locations were reprojected within each 2D markerless cameras’ Euclidean space (sagittal and frontal camera views) ^11^. Reprojected marker-based joint centre coordinates were then filtered using a bi-directional Butterworth low-pass filter (10 Hz). Once markerless and marker-based joint centre locations were filtered, event timings (heel-strike and toe-off) were obtained from Visual 3D using ground reaction force data from four imbedded force plates (Kistler 9287CA, Winterthur, Switzerland). Markerless and reprojected marker-based joint centre locations were trimmed and time normalised (101 points) to one stride (heel-strike to heel-strike) for the right and left legs separately. Left and right, ankle, knee and hip planar joint angles for both marker-based and markerless methods were calculated using the MTP, ankle, knee, hip and shoulder joint centre locations, measured in pixel units.

Statistical analysis was performed on the left and right; ankle, knee and hip angles, and MTP, ankle, knee, hip and shoulder joint centre locations in the frontal and sagittal plane, with reprojected marker-based results as the reference method ^45^. Due to the repeated measures performed with each participant (up to 10 trials, each with 101 data points), repeated measures Bland-Altman analysis where the true value varies (i.e. joint angles changes over one stride), was employed to ensure that standard deviations were not underestimated ^22^. For each joint angle and joint centre location outcome variable, this method calculates the standard deviation, and subsequently the upper and lower limits of agreement (95% confidence interval), using total variance across all data points from a one-way ANOVA (participant and residual mean square scores). In-depth detail and rationale of the repeated measures Bland-Altman analysis, with equations and examples, can be found in Supplementary File 2 and 3.

## Acknowledgements

This research was funded by CAMERA, the RCUK Centre for the Analysis of Motion, Entertainment Research and Applications, EP/M023281/1 and EP/T014865/1. We would like to thank Andrew Chapman from the Maths Resource Centre (MASH) at the University of Bath for his valuable assistance in performing the repeated measures Bland-Altman analysis.

## Author Contributions

The work was conceived by L.W., L.N., M.E., S.C., P.M., and D.C., and J.B. L.W., L.N. and M.E. collected the data. L.W., L.N. & M.E. processed the data. L.W. analysed the data and prepared manuscript and figures. All authors (L.W., L.N., M.E., S.C., P.M., D.C., and J.B.) revised and reviewed the manuscript. Funding was acquired by D.C. & J.B.

## Data Availability

Data have not been made available due to this data set being collected as part of a larger study which will be made publicly available at a later date.

