## Supplementary File 1 for "Examination of 2D frontal and sagittal markerless motion capture: Implications for 2D and 3D markerless applications"

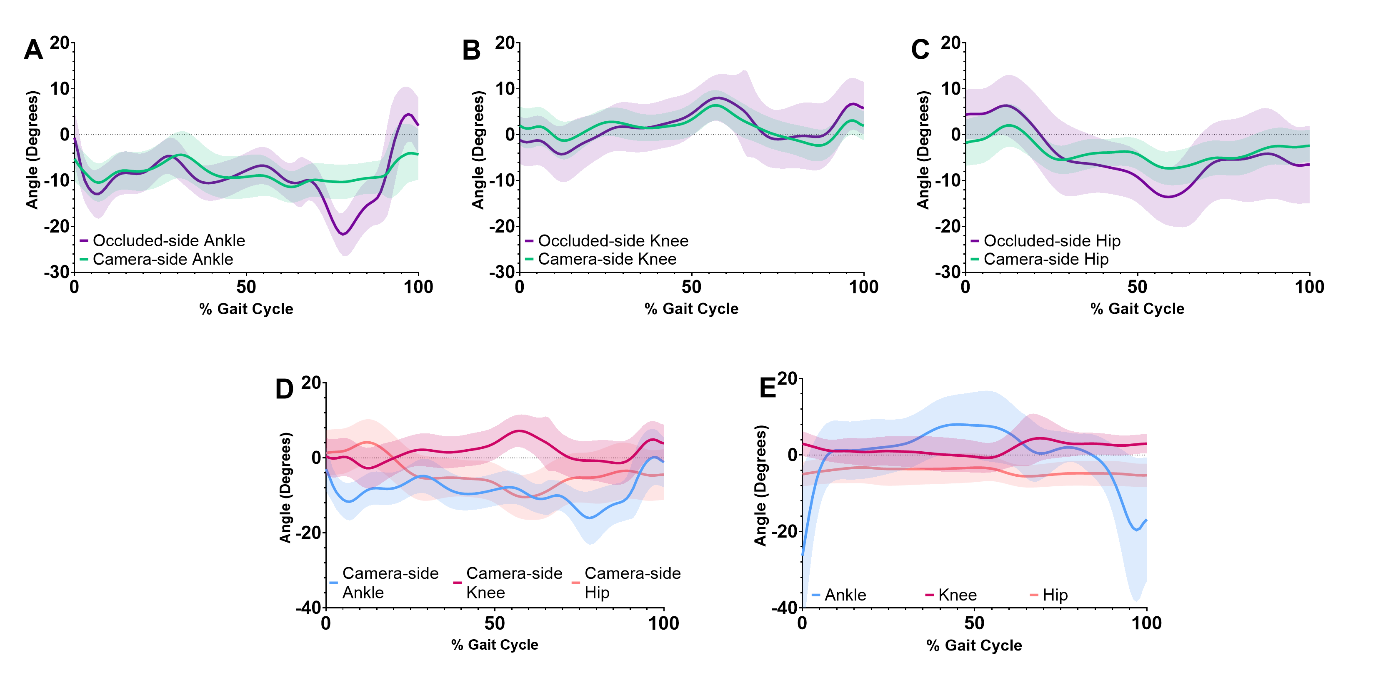

Figure 1: Markerless motion capture joint angle differences, relative to marker-based motion capture, of the ankle, knee and hip. Graph A-C compares the occluded and camera-side joints for the sagittal plane ankle (A), knee (B) and hip (C). Graph D compares the camera-side joints of the ankle, knee and hip in the sagittal plane. Graph E compares the combined left and right joint angles of the ankle, knee and hip in the frontal plane.

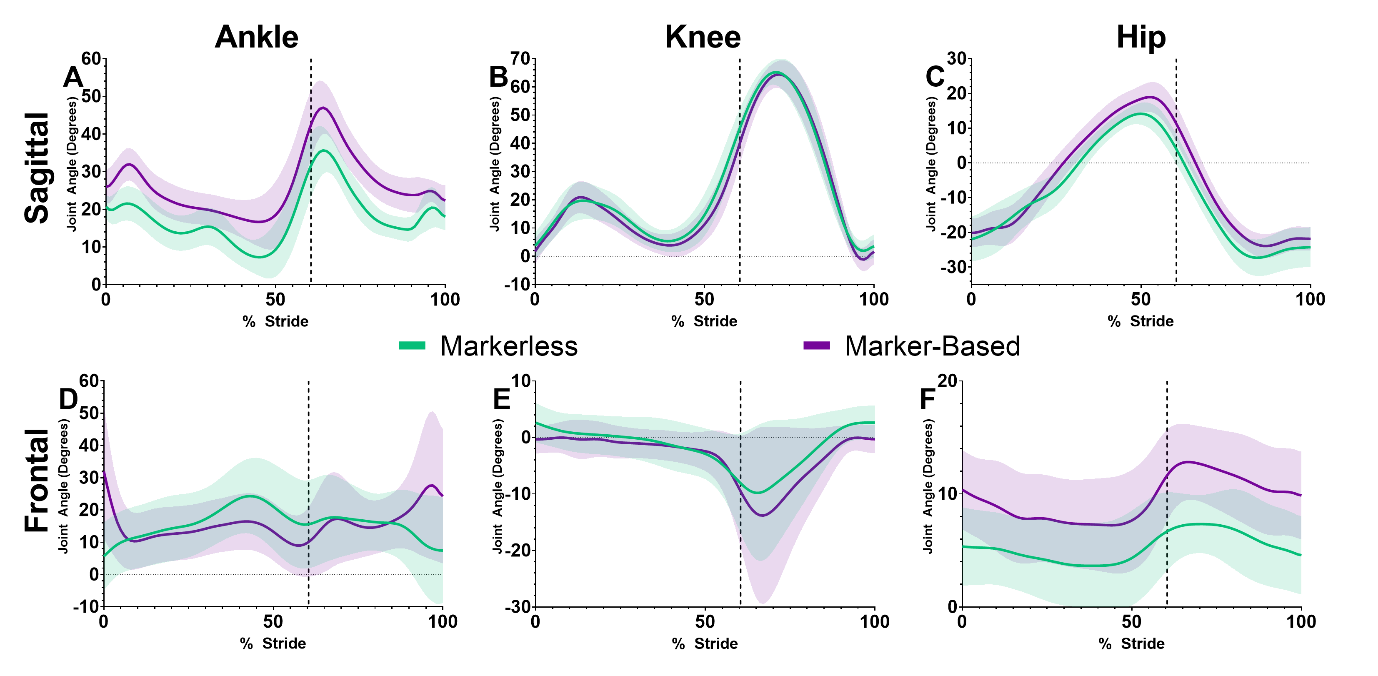

Figure 2: Comparison of joint angles (mean ± standard deviation) between markerless and marker-based motion capture in the sagittal (A,B,C) and frontal plane (D, E, F) for the ankle (A, D), knee (B, E) and hip (C, F). Sagittal plane joints are the camera-side joints only.

Table 1: Repeat measures Bland-Altman analysis of 2D markerless ankle, knee and hip joint centre locations in the sagittal and frontal plane, relative to reprojected marker-based motion capture. Camera-side represents the right side of the body which was closest to the camera, while Occluded-side represents the left side of the body which was furthest from the camera in the sagittal view. Left and right sides of the body were combined in the frontal plane.

| Joint Locations | | Bias (Pixels) | STD of Bias (Pixels) | LOA (Pixels) |
| --- | --- | --- | --- | --- |
| Sagittal Plane | | | | |
| MTP | **Camera-side** | 19.96 | 8.86 | 2.59 – 37.33 |
|  | **Occluded-side** | 23.08 | 10.35 | 2.79 – 43.37 |
| Ankle | **Camera-side** | 8.910 | 6.99 | -4.79 – 22.61 |
|  | **Occluded-side** | 8.820 | 7.46 | -5.80 – 23.44 |
| Knee | **Camera-side** | 11.95 | 6.82 | -1.42 – 25.32 |
|  | **Occluded-side** | 11.07 | 6.75 | -2.16 – 24.30 |
| Hip | **Camera-side** | 8.630 | 5.16 | -1.48 – 18.74 |
|  | **Occluded-side** | 15.22 | 6.94 | 1.62 – 28.82 |
| Shoulder | **Camera-side** | 8.51 | 3.34 | 1.96 – 15.06 |
|  | **Occluded-side** | 11.62 | 6.24 | -0.61 – 23.85 |
| Frontal Plane | | | | |
| MPT | | 8.90 | 9.61 | -9.94 – 27.74 |
| Ankle | | 7.57 | 7.23 | -6.60 – 21.74 |
| Knee | | 9.02 | 6.78 | -4.27 – 22.31 |
| Hip | | 8.90 | 6.05 | -2.96 – 20.76 |
| Shoulder | | 11.88 | 4.79 | 2.49 – 21.27 |
