## Supplementary File 2 for "Examination of 2D frontal and sagittal markerless motion capture: Implications for 2D and 3D markerless applications"

Supplementary File 2: Repeated measures Bland-Altman rationale, method and example

### Rationale

Bland-Altman analysis compares the difference between a reference method and a novel/experimental method to determine what systematic error and random error exists in the experimental method ^1^. However, a major caveat is that Bland-Altman analysis assumes all measure are independent ^1^. Biomechanical studies often record data over time, as well as over repeated trials, thus producing multiple outcome values for the same subject. To circumvent this issue, a single average measure for each participant could be obtained, and Bland-Altman analysis could then be performed on these averaged results. Unfortunately, while this application will not negatively affect the systematic bias, the standard deviation (SD) of bias will be substantially underestimated ^2^.

For example, in this study we obtained joint angle differences between marker-based motion capture and markerless motion capture, resulting in 130 trials spread across 15 participants, with each trial consisting of 101 timepoints (one stride). Therefore, there are a total of 13231 data points. As mentioned previously, one way we could analysis this dataset is to calculate the average value within each trial (average of 101 data points), reducing 13231 data points down to 130 (trials). However, because the same participants completed multiple trials, we would then be required to average these values again for each participant to meet the Bland-Altman requirement of independence, further reducing our 130 data points down to just 15 (participants). When we perform Bland-Altman analysis on these 15 data points, we obtain a mean bias of -9.3˚, a SD value of 2.5˚, with limits of agreement (LOA) of -14.2˚ to -4.4˚ (Supplementary File 3). Common-sense testing necessitates that the LOA encompass 95% of all data points. However, while this may be true for the 15 averaged values, it does not hold for the 13231 raw data points where, only 56% of these values lie within this LOA (Supplementary File 3). Thus, this approach substantially underestimates the SD and LOA. If we were to instead calculate the SD and LOA on all 130 individual trials, ignoring the fact that the same person performed multiple trials, things do not improve greatly, with a mean bias of -9.3 ˚, a SD of 2.8˚ and LOA of -14.7˚ to -3.8˚, thus only 61% of the 13231 data points fall within the 95% LOA (Supplementary File 3). As such, an adjustment to the original Bland-Altman method is needed to accurately quantify the variability (SD) between methods that are performed on repeated measures data.

In 2007, Bland and Altman published a paper entitled ‘*Agreement Between Methods of Measurement with Multiple Observations Per Individual*’ aimed to tackle this issue of repeated measures ^2^. They proposed an alternative formula for calculating the SD of repeated measures data, where the true value changes (as we have in this study) ^2^. To perform the repeated measures Bland-Altman analysis, a single dataset (table) of differences between the experimental and reference method is obtained, instead of using the raw values from both the experimental and reference method as per the original Bland-Altman method (two datasets). Supplementary File 3 depicts the table used for the example below. In the table of Supplementary File 3, columns represent each participant, and the rows encompass all time normalised data points for each participant (successful trials multiplied by 101 points). The different number of rows for each column are due to some participants having 10 successful trials (total of 1010 points) while other participants had less successful trials, due to marker dropout or other technical issues (e.g. 7 successful trials resulting in 707 data points). The mean bias value is obtained by simply taking the mean value across all data points. A one-way ANOVA is then run on this table of differences, with separate columns representing each participants data, classified as independent results (group). To calculate the SD for each joint, outputs from the ANOVA are used as inputs for the repeated measures Bland-Altman formula. The required outputs are the **between participant (between columns) mean square (MS)** and the **residual MS (within column)** values. Once SD is calculated, LOA is calculated as normal (bias + 1.96*SD).

### Method and Example

Below is a description of how to calculate the adjusted SD using the repeated measures Bland-Altman formula, with an example from our current study, using the occluded ankle joint angle difference between markerless and marker-based methods in the sagittal plane. Values have been rounded to two decimal places for ease of reading, however all decimals were retained in the workable example included in Supplementary File 3.

Table 1: Example one way ANOVA output for the occluded ankle joint angle difference between markerless and marker-based methods in the sagittal plane. Highlighted cells indicate the values used as inputs in the repeated measures Bland-Altman formula

| ANOVA | SS | DF | MS | F(DFn, DFd) |
| --- | --- | --- | --- | --- |
| Participant (between columns) | 93714 | 14 | 6694 | F (14, 13216) = 145.3 |
| Residual (within columns) | 608942 | 13216 | 46.08 |  |
| Total | 702656 | 13230 |  |  |

#### Step 0

Before beginning, we first need to calculate the **average number of observations per participant**. If the number of trials for each participant were identical, i.e. all participants had 10 trials and therefore 1010 observations, this would be as simple as taking this 1010 value. However, because we have a varying number of observations per participant, we use the equation outlined by Bland and Altman ^2^.

$$\frac{{(\Sigma m_{i})}^{2} - {\Sigma m}_{i}^{2}}{\left( N-1 \right) \times\Sigma m_{i}}=average number of observations per participant$$

Where the number of observations on participant *i*, is *m*, and *N* is equal to the total number of participants in the study.

##### Example

- *Σm_i_* = the total number of observations across all participants = 13231
- *Σm_i_^2^* = the total number of squared observations across all participants = 12292205.
- *N* = total number of participants = 15
- $\frac{{13231}^{2} - 12292205}{\left( 15-1 \right) \times13231}= 878.71$
- Thus, the number of observations per participant in this study was 878.71.

#### Step 1

From Table 1, we take the **Participant MS (between columns)** value and then minus the **Residual MS (within columns),** to calculate the **average difference across participants**.

##### Example

- $6694 - 46.08 = 6647.92$

#### Step 2

We then divide the **average difference across participants** (Step 1), by the **number of observations per participant** (Step 0), to calculate the **estimated component of variance**, which represents the heterogeneity across all participants.

##### Example

- $\frac{6647.92}{878.71}=7.56$

#### Step 3

The **estimated difference across participants** (Step 2) is then added to the **Residual MS values (within column)** from Table 1, to calculate the **total variance**.

##### Example

- $7.56+46.08=53.64$

Step 4: Finally, the square root of the **total variance** (Step 3) is calculated to determine the **standard deviation**.

##### Example

- $\sqrt{53.64}=7.32$

Thus in our example, we obtained a mean bias of -9.3 ˚, a SD of 7.32˚, with LOA of -23.66˚ to 5.05˚, which common sense testing indicated that 93.9% of all 13231 datapoints fell within this range (Supplementary File 3). Across all sagittal and frontal joint angle measures, common sense testing averaged out to 95.5% of all individual datapoints falling within the range of the repeated measure 95% LOA. Therefore, the repeated measures Bland-Altman analysis appears to work well for this application.

It should be noted that the repeated measures Bland-Altman method treats the differences within a participant (rows) as independent points. By their own admission, Bland and Altman ^2^ admit this is a large assumption, however they concluded that this was unlikely to have a major impact on their results. To account for the fact that timepoints are not independent, Myles and Cui ^3^ used linear mixed modelling to obtain MS scores that account for changes over time. However, linear mixed modelling assumes that any change over time would be linear. For our application, the difference between the marker-based and markerless methods over one stride were non-linear (Supplementary File 1). As such, employing a linear mixed model to account for these changes would introduce additional error into the model. Therefore, we arrived at a similar conclusion to Bland and Altman ^2^, in that treating these within participant datapoints as independent is a suitable approach which is supported by our common sense testing.
